# Electromigration of Charged Analytes Through Immiscible Fluids in Multiphasic Electrophoresis

**DOI:** 10.1101/2024.05.29.596534

**Authors:** Md Nazibul Islam, Yang Liu, Amy E Herr

**Affiliations:** Department of Bioengineering, University of California, Berkeley, CA 94720, USA; School of Chemical, Materials and Biomedical Engineering, University of Georgia, Athens, GA 30602, USA; Chan Zuckerberg Biohub, San Francisco, CA 94158

## Abstract

Multiphasic buffer systems have been of greatest interest in electrophoresis and liquid-liquid electrotransfer; this study extends that foundation by exploring the interplay of the geometric and viscous properties of an interleaving oil layer on the electrotransfer of a charged analyte from an aqueous solution into a hydrogel. We utilized finite element analysis to examine two complementary configurations: one being electrotransfer of a charged analyte (protein) in an aqueous phase into a surrounding hydrogel layer and another being electrotransfer of the protein from that originating aqueous phase – through an interleaving oil layer of predetermined viscosity and thickness – and into a surrounding hydrogel layer. Results indicate that the presence of an oil layer leads to increased skew of the injected peak. To explain this difference in injection dispersion, we utilize Probstein’s framework and compare the Péclet (Pe) number with the ratio between length scales characteristic to the axial and radial dispersion, respectively. The formulation assigns electrotransfer conditions into six different dispersion regimes. We show that the presence or absence of an interleaving oil layer moves the observed peak dispersion into distinct electrotransfer regimes; the presence of an oil layer augments the electrophoretic mobility mismatch between the different phases, resulting in a five-fold increase in Pe and a six-fold increase in the ratio between the axial to radial dispersion characteristic lengths. We further show that oil viscosity significantly influences resultant injection dispersion. A decrease in oil-layer viscosity from 0.08 Pa·s to 0.02 Pa·s results in a >100% decrease in injection dispersion. Our theoretical predictions were experimentally validated by comparing the electrotransfer regimes of three different mineral oil samples. We show that lowering the oil viscosity to 0.0039 Pa·s results in an injection regime similar to that of the absence of an oil layer. Additionally, we measure the migration distance and show that average electromigration velocity over the transit duration is inversely proportional to the viscosity of an interleaving oil layer. Understanding of the impact of electrotransfer of charged species across multiple immiscible fluid layers on peak dispersion informs the design of multiphasic electrophoresis systems.

## 1. Introduction

Electric field-induced mass transfer (electrotransfer) of charged materials across liquid-liquid interfaces plays an important role in developing chemical and biomedical systems, including product extraction, target separation, microextraction across droplets, and the development of electrofluidic displays ^1–12^. Over the years, researchers have made significant advances in enhancing our understanding of electrotransfer mechanisms. In a foundational study by Austin et al. (1971), the authors demonstrated a reduction in effective interfacial tension at an electrically charged liquid-liquid interface, leading to interfacial-tension-induced surface flows and increased mass transfer rates ^13^. Later, Mohan et al. (1983) confirmed that the application of an electric field across a liquid-liquid interface causes Marangoni-type instabilities due to different susceptibilities of the two phases to the field. These instabilities and the resulting mixing enhance the rates of mass transfer across the interface ^14^. In 1993, He and Baird demonstrated a fourfold increase in the mass transfer of benzoic acid from water droplets to a continuous phase of mineral oil in the presence of an applied electric field. The authors demonstrated a reduction in droplet diameter and an increase in droplet initial velocity, thus an overall increase in interfacial area in the presence of the electric field leads to higher mass transfer rates ^15^. The authors expanded upon this earlier work by exploring pulsed electric fields and in 1997, showed that pulsed electric fields are more energy-efficient in terms of power consumption than DC electric fields, although the latter generates higher mass transfer rates ^16^. In 2002, Kornyshev et al. explored the dynamics of ion transfer across the interface of two immiscible electrolyte solutions (ITIES). The authors demonstrated that a favorable electric field orientation enhances the protrusion at the interface in a way that facilitates quicker ion transfer by effectively “pulling” the ion through the interface with reduced resistance ^17^. More recently, Weatherley et al. reported a detailed study on the formation of electrically induced disturbances and enhancement of mass transfer and reaction rates in liquid-liquid systems. The authors studied three different systems: a pendant droplet of aqueous ethanol in a continuous n-decanol phase, a planar interface constituting the same set of liquids, and an enzymatic hydrolysis system developed by electrostatic spraying of aqueous enzyme solutions into an immiscible oil phase. The authors theoretically and experimentally demonstrated an increase in interfacial disturbance in the presence of the applied electric field, which increased mass transfer in all three systems and reaction rates in the third system ^18^.

Based on these theoretical understandings, several multiphasic electrophoresis applications have been recently reported. For example, Gan et al. demonstrated protonation-induced molecular permeation of red ink molecules containing a single amine group from water to oil under an applied electric field ^12^. This understanding led to the development of an efficient electrofluidic display ^19^. Further applications have extended into the biomedical field. For instance, Abdelgawad et al. demonstrated sample pre-processing and chemical separation on a single chip by developing a hybrid microfluidic system where an electric field was applied to drive the contents of droplets containing samples such as fluorescent dyes, amino acids, and cell lysate from the digital microfluidic platform into microchannels through electrokinetic flow ^20^. Kee et al. have developed an aqueous two-phase (ATP) electrophoresis system to recover and purify extracellular keratinase from *Kytococcus sedentarius* TWHKC01 fermentation broth. The researchers reported a two-fold increase in selectivity by the introduction of electric field ^11^. Sauer et al. demonstrated that electric-field-assisted packed-bed milli-reactors can be effectively used to control reaction conversion and enhance the enantiomeric ratio in the production of pharmaceutical intermediates ^10^. Petersen et al. demonstrated the electroextraction of basic drugs such as Pethidine, Nortriptyline, Methadone, Haloperidol, and Loperamide from a donor droplet sample consisting of biological fluids such as human urine and plasma (10 μL) into an acceptor reservoir ^9^. Additionally, there has been a recent uptake in the development of electric field-assisted extraction and separation using microfluidic ATP systems in both continuous and batch phases. These techniques have been used to separate size-based DNA strands, identify, and purify cells, extract amino acids, separate proteins, extract cocaine and lidocaine in artificial saliva, malachite green in tap water, and bisphenol A (BPA) in red wine and extract crystal violet (CV) and leuco crystal violet (LCV) from fish muscle ^1–7^.

Building on these advancements, recently, our lab has developed a hybrid microfluidic system termed “DropBlot”, which combines water-in-oil droplet-based single-cell sample preparation with microfluidic and western blotting technologies ^21^. This platform aims to facilitate the understanding of putative protein biomarkers using archived cell and tissue specimens. In the DropBlot workflow, we encapsulate paraformaldehyde-fixed and methanol-fixed cells in aqueous droplets containing a lysis buffer in a continuous mineral oil phase, lyse the cells, and then allow the droplets to gravity-settle in a planar polyacrylamide microwell array. We then perform single-cell western blot analysis, a technique introduced by our lab ^22,23^, measure a priori identified protein targets. DropBlot enables precise protein biomarker discovery at a single-cell resolution, benefiting research where assessing cellular heterogeneity is important ^24^.

Complementing that translational biomedical study, here we seek to understand injection dispersion arising from electromigration of solubilized protein contained in the aqueous core of a water-in-oil droplet through a thin-oil layer and into the molecular sieving gel of polyacrylamide gel electrophoresis (PAGE). Since reducing injection dispersion enhances the separation resolution and detection sensitivity of PAGE, we seek to understand the dispersion source and how to mitigate any resultant injection dispersion in this multiphasic electrophoresis system.

## 2. Materials and Methods

### 2.1 Finite Element (FE) Analysis

We have developed a 2-D FE model using a commercially available FE Multiphysics package, COMSOL Inc. (Burlington, MA). This model comprises 2,872 triangular mesh elements, with the maximum and minimum element sizes being 2 μm and 0.01 μm, respectively. The average element quality of our model, evaluated based on the skewness of the triangular mesh elements (with 1 representing the ideal equilateral triangular mesh), is 0.87. This indicates that the shape of the mesh elements closely approximates the ideal configuration. Utilizing the electric current and diluted species transport modules, we analyzed protein movement over 40 s under an externally applied electric field of 40 V/cm for the two scenarios, as described in the results and discussion section.

### 2.2 Mineral Oil Electrical Conductivity

The electrical conductivity of the mineral oil sample (ASI Standards, Baton Rouge, LA) was measured using a Keithley 2410 Sourcemeter, and the results are presented in the table below:

| Sample | Electrical Conductivity (S/m) |
| --- | --- |
| Mineral Oil | Below detection limit (BDL) |
| Mineral Oil + 2% (v/v) Span 80 | 8.14X10<sup>-9</sup> |

### 2.3 Design and Fabrication of Single-emulsion Droplet-generation Chip

In order to generate aqueous droplets containing protein, we designed the single-emulsion droplet generation devices using AutoCAD 2024 (Autodesk, San Francisco, CA). We utilized soft lithography for fabricating the silicone masters. We used SU-8 3050 (Kayaku Advanced Materials, Westborough, MA), with a target height of 60μm, following the guidelines provided by the manufacturer. The polydimethylsiloxane (PDMS) was prepared using a Sylgard 184 Silicone Elastomer Kit (Ellsworth Adhesives, Germantown, WI), mixing at a ratio of 10:1. After curing the mixture at 70ºC for 3 hr, we baked the PDMS devices for an additional 48 hr at 80ºC to restore the hydrophobic characteristics.

### 2.4 Design and Fabrication of Open-fluidic Single-cell Western Blotting Chip

To perform western blot on the contents of a single droplet, we fabricated a polyacrylamide gel western blotting clip composed of planar arrays of microwells, each abutting a polyacrylamide gel electrophoresis regions. We used our previously published methodology for wafer microfabrication and silanization ^22^. The microwell array comprises 5,000 microwells, each with a diameter of 50 μm and a depth of 60 μm. The spacing between microwells is 1 mm in the electrophoresis direction and 300 μm perpendicular to electrophoresis. To prepare an 8%T polyacrylamide gel, we mixed the gel precursor solution with 30% (w/w) acrylamide/bis-acrylamide (Sigma-Aldrich, St. Louis, MO), N- (3-((3-benzoylphenyl)formamido)propyl) methacrylamide (BPMA, PharmAgra Labs, Brevard, NC), 10× Tris-glycine buffer (Sigma-Aldrich, St. Louis, MO), and ddH_2_O (Sigma-Aldrich, St. Louis, MO). We initiated and maintained the chemical polymerization of gels for 20 minutes using 0.08% (w/v) ammonium persulfate (APS, Sigma-Aldrich, St. Louis, MO) and 0.08% (v/v) TEMED (Sigma-Aldrich, St. Louis, MO). Following polymerization, we carefully removed the gels from the wafers through a delamination method using a razor blade. The gels were then stored in deionized (DI) water for future use.

### 2.5 Integration of Droplets with Micro-Electrophoresis Array

To study protein electrotransfer from droplets through a thin oil layer and into a polyacrylamide gel, we diluted fluorescently tagged BSA protein (Sigma-Aldrich, St. Louis, MO) to a concentration of 0.1 mg/mL and encapsulated the solution in 45-μm droplets using the single-emulsion droplet-generation chip. Mineral oil samples with three different viscosities (0.0039, 0.022 and 0.065 Pa·s) (ASI Standards, Baton Rouge, LA) were used as continuous phase. These protein-laden droplets were gravity-settled onto the surface of the open-fluidic single-cell western blotting chip containing PA gel. Once the droplets settled in the microwells, we sealed the gel with a hydrophobic glass slide and gently washed the gel with DI water to remove excess oil and hydrate the hydrogel. This assembly was then placed in a custom-designed electrophoresis chamber consisting of a plastic chamber with two platinum electrodes at opposite ends of the chamber. A 12.5 mL aliquot of lysis buffer (1× Tris-glycine and 0.5% (v/v) sodium dodecyl sulfate (SDS, Sigma-Aldrich, St. Louis, MO) was poured into the chamber and allowed to equilibrate for 20 s. For PAGE, a constant electric field of 40 V/cm was applied using a DC power supply (Bio-Rad PowerPac Basic, Hercules, CA) for 40 s. Upon the completion of PAGE, the applied voltage was set to floating, and the protein bands were photo-captured into the PA gel through a 45 s UV exposure (Hamamatsu Lightingcure LC5 UV source, Hamamatsu Photonics, Japan) that activates hydrogen abstraction and subsequent covalent attachment of protein species to benzophenone in the polymer. The chips were rinsed with deionized (DI) water and then dried with nitrogen gas prior to imaging with an inverted fluorescence microscope and a Genepix® microarray scanner (4300A, Molecular Devices, San Jose, CA).

## 3. Results and Discussion

Our study aimed to explore the effect electrotransfer of a charged analyte (protein) from the aqueous phase of a water-in-oil droplet, through the oil layer of said droplet and into a supporting polyacrylamide gel that acts as a molecular-sieving matrix for terminal polyacrylamide gel electrophoresis (PAGE), (Figure 1). The ultimate goal of the study is the development of design guidelines for multiphasic electrophoresis systems such as DropBlot, ATP electrophoresis, and protonation-induced molecular permeation.

**Figure 1:**
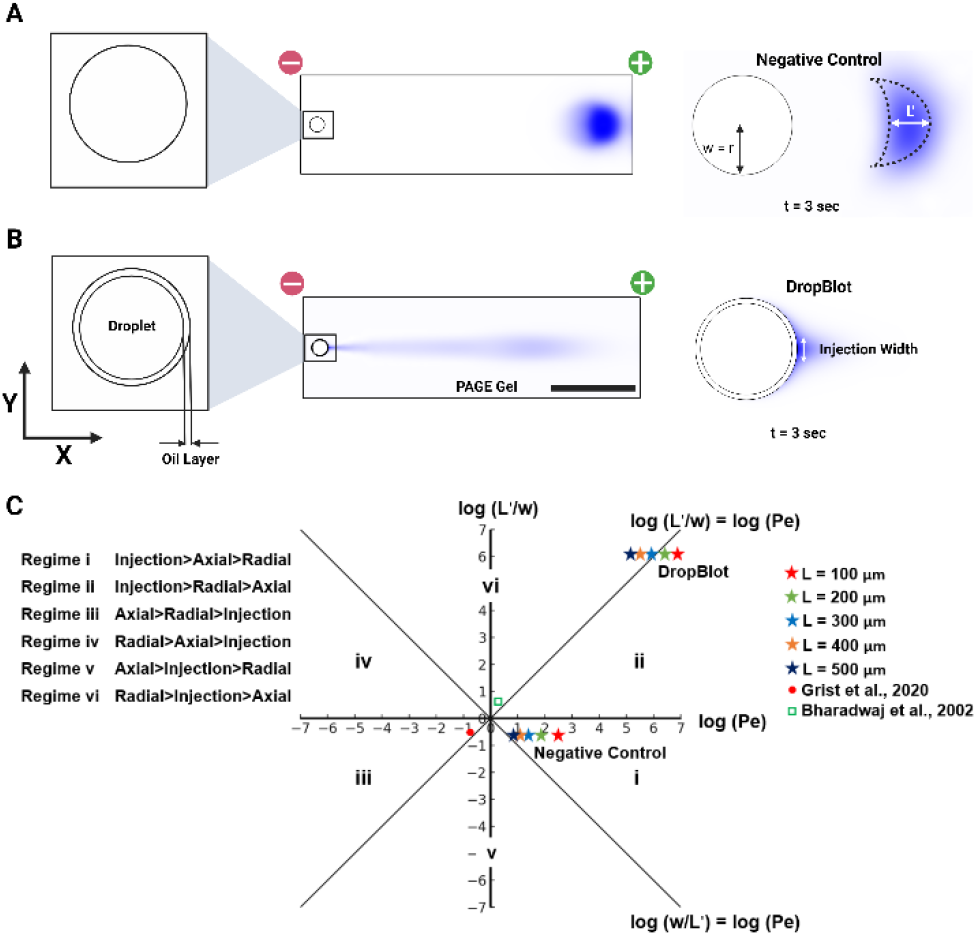
Finite element model and Péclet regime analysis reveal that protein electrotransfer dynamics in various open fluidic devices are determined by device configuration and Péclet regimes. (A) Numerical simulation of protein electrotransfer from a microwell into a polyacrylamide gel shows effective protein electrophoretic mobility (scale bar: 250 μm). (B) Numerical simulation highlights how a thin oil layer modifies electrotransfer from a droplet into a polyacrylamide gel, resulting in a narrower injection width and a longer injection tail. (C) Péclet regimes explain variations in injection shapes, with increased Péclet numbers and axial to radial diffusion length ratios in the presence of an oil layer compared to negative control, due to higher disparities in protein mobilities between the oil and gel phases.

We commence our investigation by comparing two complementary electrotransfer configurations: 1) a 50-micron microwell containing a 2-μM Bovine Serum Albumin (BSA) solution as a negative control and 2) a 45-micron water-in-oil droplet encapsulating the same concentration of BSA in a 2.5-micron thick mineral oil layer to mimic protein electrotransfer in a DropBlot assay. We use BSA as a model protein as we expect BSA to be representative of most proteins with respect to the relative electrophoretic mobility in the droplet (aqueous), oil layer, and PAGE gel phases. By employing FE analysis, we developed a model for estimating protein dispersion in each configuration (Figure 1A, B).

In considering electrotransfer of a charged analyte from an aqueous starting phase through a thin oil layer and into a hydrogel molecular sieving matrix, we asked: what distinguishes the regimes of dispersive behavior under these conditions? To tackle this question, we devised a non-dimensionalization approach inspired by Probstein’s methodology ^25^, enabling us to compare the thin oil layer’s impact with that observed in the negative control. We introduce L’ and w as the axial and radial dispersion characteristic length scales and compare the ratio between L’/w and Peclet number (Pe) (see SI) ^26^.

A Pe>L’/w indicates that t_radial_ and t_axial_ >> t_injection_ and therefore the injection shape will have a greater impact on the injection dispersion than either radial or axial diffusion.

Conversely, Pe<L’/w indicates that axial and radial diffusion dominate injection dispersion over injection shape ^26–29^.

By comparing Pe and L’/w with unity we can divide the conditions into six different regimes (Figure 1C). We compare the negative control with the configuration including an oil layer across various separation lengths (100 to 500 μm) to determine the dominant dispersion source for each scenario. For the configuration with an oil layer, the injection width for calculating w was determined from the FE simulation (Figure 1B). Using equations 1 and 2 (SI), we calculated L’/w and Pe, and we observed both log(L’/w) and log(Pe) are greater than 1. This is because L’ is dependent on the ratio of (μ_ii_/μ_i_). In the case of the oil layer, μ_ii_ is the electrophoretic mobility of BSA in gel and μ_i_ is the electrophoretic mobility in oil. Since μ_ii_ >> μ_i_, therefore L’>w and a higher value of Pe (1.9×10^5^ to 6.8×10^6^). This higher mismatch in electrophoretic mobility between the oil phase and gel phase results in a higher injection and radial variance (SI, Equations 4 and 5) which results in a higher overall dispersion. We observe that in low separation distance (L<300 μm) Pe>L’/w which indicates injection dominated dispersion whereas at higher L, we observe L’/w>Pe which indicates radial diffusion dominated dispersion. When we compare this data with negative control, we observe a lower value of Pe (7.6 to 309), and L’/w is less than 1. The reason is that the electrophoretic mobility of BSA in free solution is greater than that in the gel, therefore L’<w (for negative control w: microwell radius). These values are in accordance with previously published results for scPAGE ^26^. We also observed a sharp increase in local electric field strength within the oil layer, while the electric field strength remains constant in other parts of the system; notably, it remains uniform throughout the separation lane (SI, figure S1). This sharp increase in the oil layer is attributable to the significantly low electrical conductivity of mineral oil. The local field strength must increase to maintain continuity across the different phases of the system.

In addition, we observe a shift from injection-dominated dispersion to axial dominated dispersion as we gradually increase the separation length. The relatively low value of L’/w and Pe is a result of a lower difference in electrophoretic mobility between free solution and PAGE gel which results in lower dispersion variance compared to the scenario with an oil layer between the free solution and the gel. In addition, we compared our findings with those from two previously published electrokinetic injection systems: 3D projection electrophoresis and electrokinetic T-injection. For single-cell 3D projection electrophoresis, our observations indicate that both radial and axial diffusion significantly influence dispersion at a separation length of 1mm ^30^. Conversely, in the case of electrokinetic T-injection, we identified Taylor-Aris dispersion as the predominant mechanism ^31^.

Drawing on the insights from Probstein’s methodology, we aimed to devise strategies to minimize the radial and injection dispersion observed in droplet-oil-gel systems. Previous experimental findings have shown that a reduction in oil layer thickness can reduce protein injection dispersion. Therefore, the first strategy we investigated involved reducing the thickness of the oil layer to observe its impact on the dispersion of protein injection. To decrease the oil layer thickness while keeping the microwell cross-section constant, we used elliptical microwells to facilitate droplet settling into the microwell. Using Poisson distribution, we analyzed droplet settling into microwells of varying eccentricities and found that microwell eccentricity does not affect droplet settling for the analyzed range (0 to 0.57) (SI, figure S2).

We employed FE simulations to investigate protein injection dispersion across three different oil layer thicknesses. Our initial thickness was set at 2.5 μm, corresponding to the original DropBlot assay, and was subsequently decreased in 1 μm increments. We then compared the results to a scenario with zero oil layer thickness (Figure 2A). Using the method reported by Liu and Herr ^21^, we calculated and plotted the dispersion in the gel phase along the x-axis and y-axis for each oil layer thickness and noted a progressive reduction in x-axis variance with decreasing oil layer thickness (y-axis variance is negligible compared to x-axis variance) (Figure 2B). To further explore the underlying dispersion mechanism, we applied Probstein’s method, as illustrated in Figure 2C (L: 1 mm). This point is crucial as we encounter two distinct interfaces in this context: 1) droplet to oil, and 2) oil to gel, resulting in two different sets of Peclet numbers (Pe) and L’/w ratios for each oil layer thickness. In our FE analysis, we focused exclusively on the results pertaining to the gel phase, as the simulation of the oil phase necessitated molecular-level modeling that was beyond the scope of this study ^17^.

**Figure 2:**
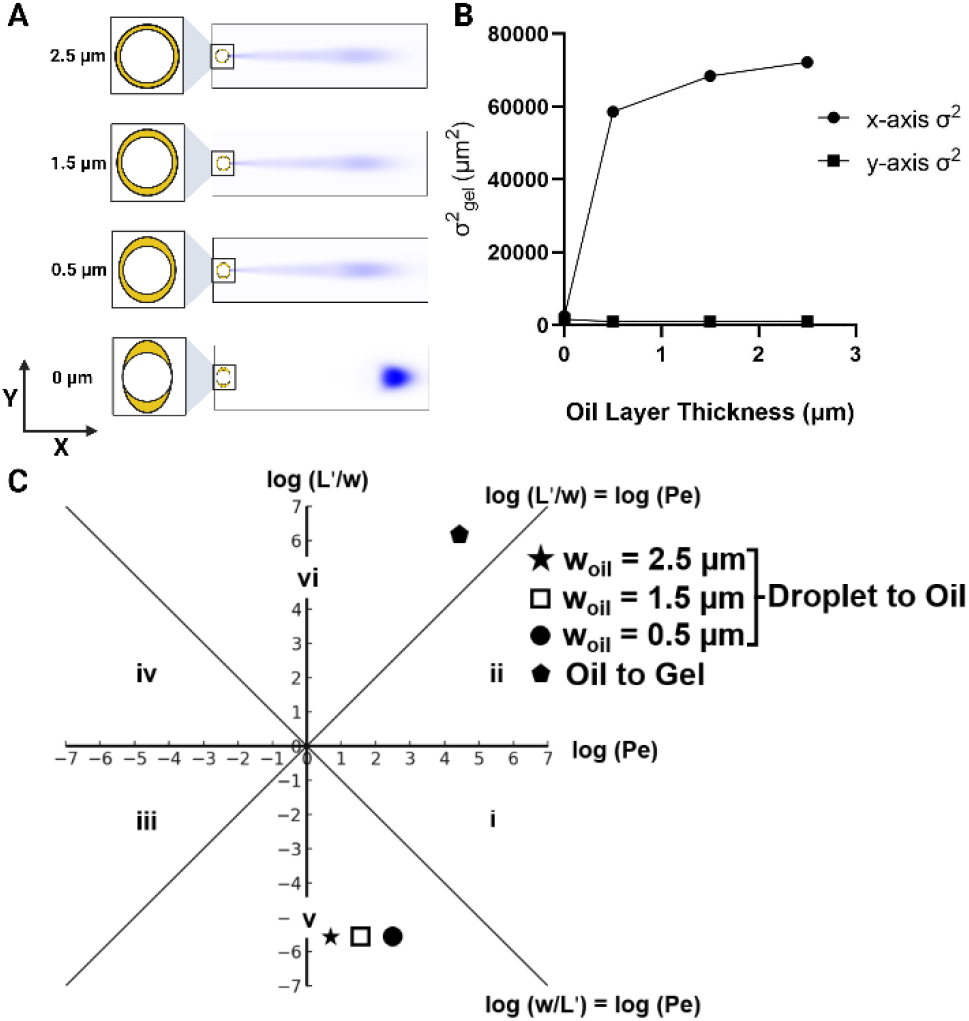
Relationship between protein electrotransfer and oil layer thickness demonstrates minimal impact on layer variability. (A) Numerical simulation of protein electrotransfer across different oil layer thicknesses indicates that changes in oil layer thickness have a negligible effect on electrotransfer compared to the absence of an oil layer. (B) X and y-axis variance versus oil layer thickness as a measure of dispersion, with the variance along the y-axis being negligible compared to that along the x-axis. (C) Péclet regime analysis for the droplet-to-oil and oil-to-gel interfaces across varying oil layer thicknesses, indicating that a decrease in oil layer thickness slightly reduces the variance in protein injection by increasing the Péclet number (Pe).

For droplet to oil interface, we observe L’/w<1. This is because the mobility of protein in a droplet is significantly higher than in the oil, leading to L’<w. Reducing the separation length (L) from 2.5 μm to 0.5 μm resulted in an increase in the Pe number from 5.9 to 249, yet the L’/w ratio remained unaffected, as changes in L do not influence this ratio. Conversely, as the protein transitions from oil to gel, both L’/w and Pe >>1, as discussed previously. Adjustments in the oil layer thickness did not alter the Pe and L’/w values at the oil-to-gel interface. Therefore, the observed 20% reduction in dispersion upon reducing the oil layer thickness from 2.5 μm to 0.5 μm can be attributed to the enhanced electrophoretic advection of the protein within the oil layer (increase in Pe). Nevertheless, compared to the scenario with zero oil layer thickness (187% reduction compared to the 2.5 μm oil layer thickness), this reduction in injection dispersion is marginal.

Since the diffusivity and, consequently, the electrophoretic mobility of a protein is influenced by the viscosity of the medium, we aimed to determine how changes in oil viscosity would affect protein injection dispersion. We utilized our FE model to analyze injection dispersion across four different oil viscosities, starting from 0.08 Pa·s, which is closer to mineral oil, and down to 0.02 Pa·s, which is closer to water with 0.02 Pa·s increments. We then plotted the resulting gel phase dispersion variance along both the x-axis and y-axis. As illustrated in Figures 3A and B, a 133% reduction in x-axis variance occurs with decreasing oil viscosity from 0.08 Pa·s to 0.02 Pa·s (y-axis variance negligible compared to x-axis variance). Applying Probstein’s framework we noted an increase in both the Peclet number (Pe) from 8.32 to 204 and the L’/w ratio at the droplet to oil interface, and a decrease in these parameters (Pe: 24,000 to 1,000) at the oil to gel interface as we lower the oil viscosity (L: 1 mm) (Figure 3C). This phenomenon occurs because a reduction in oil viscosity diminishes the electrophoretic mobility mismatch between the droplet, oil, and gel phases (viscosity is inversely proportional to electrophoretic mobility). Consequently, the decreased electrophoretic mobility mismatch at both interfaces lessens radial and injection dispersion (Equations 4 and 5), leading to a diminished overall dispersion. A progressive reduction in oil viscosity (from 0.1 Pa·s to 0.001 Pa·s, equating to water’s viscosity) shifts the system from Figure 1B towards Figure 1A, thereby reducing dispersion variance.

**Figure 3:**
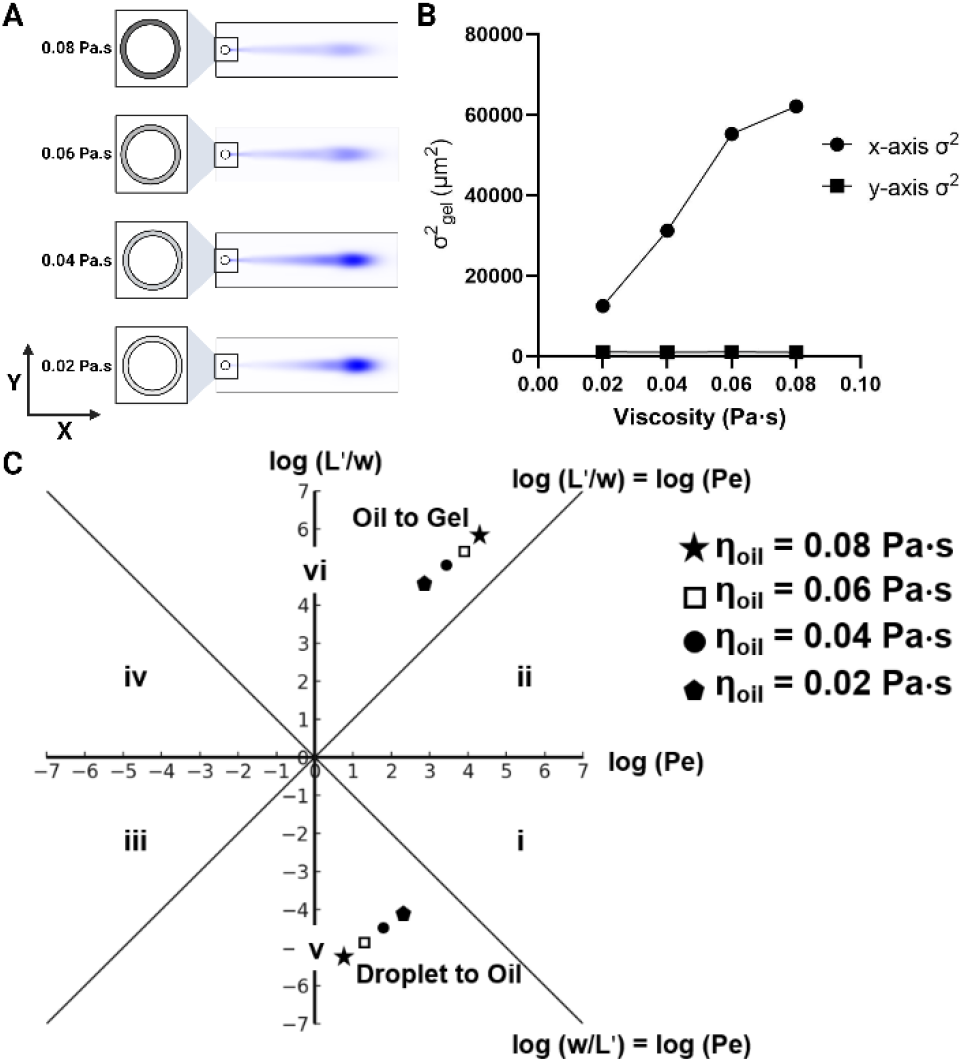
Relationship between protein electrotransfer and oil layer viscosity demonstrates a decrease in oil viscosity reduces injection variance. (A) Numerical simulation of protein electrotransfer across oils of varying viscosities, showing that a decrease in oil viscosity facilitates improved protein migration through the thin oil layer. (B) X and y-axis variance versus oil layer viscosity, with the variance along the y-axis being negligible in comparison to the x-axis. (C) Péclet regime analysis for the droplet-to-oil and oil-to-gel interfaces across different oil viscosities. A decrease in oil viscosity leads to an increase in both Pe and (L’/w) at the droplet to oil interface and a decrease at the oil to gel interface, thus minimizing the variance in protein migration between the droplet to oil and oil to gel phases.

Leveraging our theoretical understanding, we next experimentally scrutinized injection dispersion resulting from charged analyte electrotransfer through one of three mineral oil layers, each layer presenting a different viscosity oil.

In our study, we aimed to observe protein injection dispersion through a thin oil layer, using mineral oil samples with three different viscosities (0.0039, 0.022 and 0.065 Pa·s). We assumed negligible change in oil viscosity of the oil due to an increase in temperature from Joule heating, as the thermal conductivity of mineral oil is five times lower than that of water^32^. Our observations indicate protein retention at the oil-gel interface thus resulting in two distinct peaks during intensity measurement in the x-direction. Therefore, for experimental observation, we report protein migration distance instead of injection dispersion and employ Probstein’s framework to compare between experimental and theoretical findings. Our results demonstrate that a reduction in oil viscosity resulted in larger protein migration distances (Figure 4A), with a significant statistical difference in migration distances between the 0.0039 Pa·s oil and the more viscous oil samples (Figure 4B). Applying Probstein’s framework, we noted that for 0.0039 Pa·s oil viscosity, electrotransfer from the droplet to the oil transitioned to the same regime as the negative control. For the electrotransfer from oil to gel, we observed a decrease in both Pe and L’/w ratios due to the reduction in electrophoretic mobility mismatch (Figure 4C). These experimental observations align with our theoretical predictions. Therefore, based on the simulation and experimental data, we establish the following design rule: when performing electrotransfer across multiphasic systems, the Log (Pe) vs Log (L’/w) among the different phases should converge in order to reduce dispersion and increase migration distance. This can be achieved by lowering the viscosity mismatch between the immiscible fluids.

**Figure 4:**
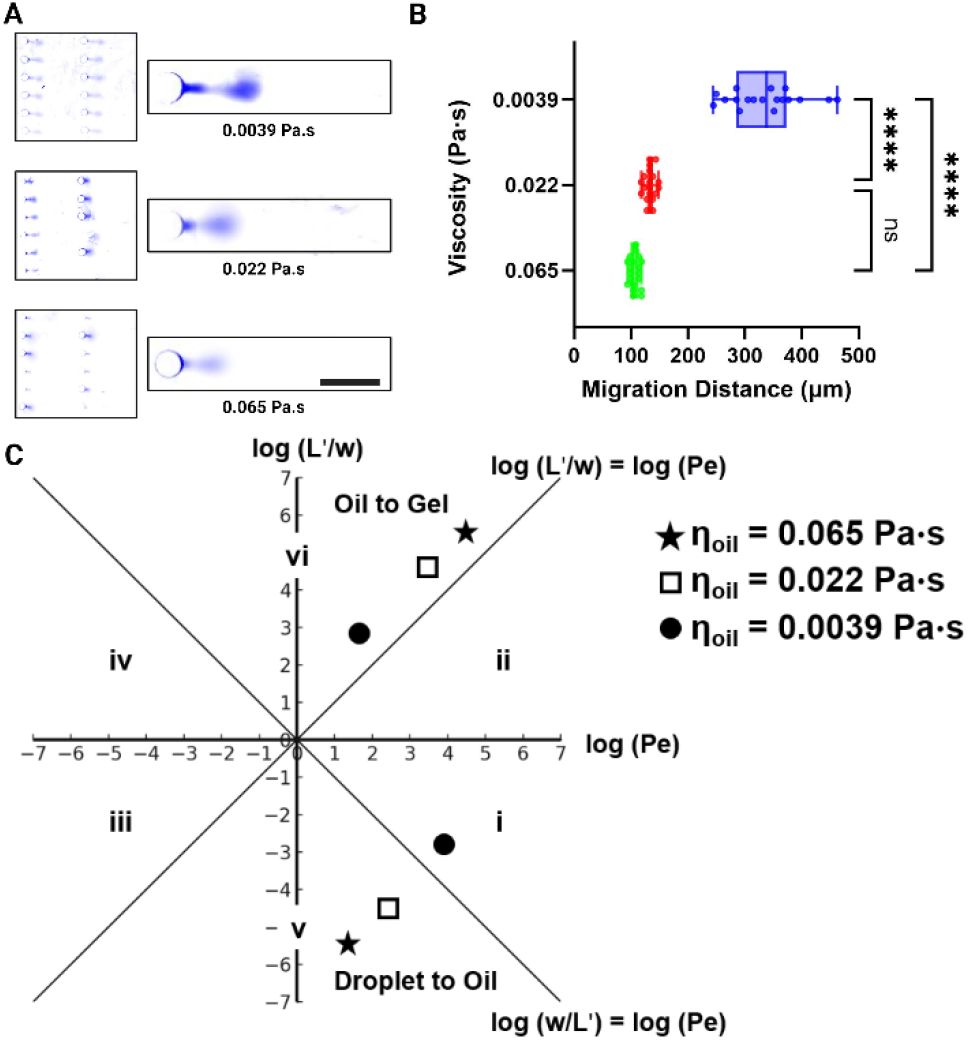
Empirical results of electrophoretic migration of BSA from aqueous droplets into polyacrylamide gel through a thin oil layer of varying viscosity. (A) Micrographs of protein migration for three different oil viscosities, a decrease in protein migration distance is observed with an increase in mineral oil viscosity. The increase in mineral oil viscosity leads to an increased electrophoretic mobility mismatch among the aqueous, oil, and gel phases, which, in turn, reduces the migration distance (scale bar: 250 μm). (B) Migration distance vs oil layer viscosity, a statistically significant difference in migration distance is observed between the 0.0039 Pa·s oil and more viscous oil samples. (C) Péclet regime analysis for the droplet-to-oil and oil-to-gel interfaces across different experimental conditions. A decrease in oil viscosity decreases electrophoretic mobility mismatch between aqueous, oil and gel phases thus increasing migration distance.

## 4. Conclusion

In this study, we added a design guideline for multiphasic electrophoresis systems by examining the electrophoretic injection of a protein sample through a thin oil layer into a polyacrylamide gel. We developed an FE analysis model and employed a non-dimensionalization technique based on Probstein’s methodology to explore the impact of the oil layer on the protein injection profile. Our findings indicated that the presence of the oil layer increases the Peclet number (from 10^2^ to 10^6^) and the ratio between characteristic length scales, thereby enhancing dispersion. To mitigate injection dispersion, we investigated the effects of varying the oil layer’s thickness and viscosity. Our numerical analysis suggested that altering the thickness of the oil layer has a minimal impact on reducing injection dispersion. In contrast, changing the oil viscosity has a more substantial effect, with a reduction in oil viscosity leading to a decrease in injection dispersion. This reduction is attributed to the decreased electrophoretic mobility mismatch between the different phases, which in turn reduces interfacial stacking. We experimentally validated our theoretical findings using three different mineral oil samples with varying viscosities. The experimental results confirmed that protein migration distance is significantly higher when using a mineral oil with a viscosity of 0.0039 Pa·s compared to oils with viscosities of 0.022 and 0.065 Pa·s. These findings will be useful in designing multiphasic electrophoresis systems by manipulating medium viscosity mismatch to separate or concentrate a target of interest.

## ASSOCIATED CONTENT

### Supporting Information

The Supporting Information is available free of charge on the ACS Publications website.

S.I (file type, i.e., PDF)

## AUTHOR INFORMATION

### Corresponding Author

* Amy E Herr, Department of Bioengineering, University of California, Berkeley, CA 94720, USA & Chan Zuckerberg Biohub, San Francisco, CA 94158.

### Author Contributions

M.N.I and A.E.H designed the experimental and theoretical framework. M.N.I performed the finite element analysis, non-dimensionalization analysis and experiments. Y.L and A.E.H designed the hybrid microfluidic platform. Y.L developed the droplet loading protocol. All authors have given approval to the final version of the manuscript.

## ACKNOWLEDGMENT

This work was funded by the Chan Zuckerberg Biohub Investigators Program and by the National Institutes of Health (NIH) R01CA20301. The photolithography was performed in the QB3 Biomolecular Nanotechnology Center at UC Berkeley. The figures were created with BioRender.com. We sincerely thank Herr lab member Maya Overton for initial training on DropBlot assay. We acknowledge all lab members of the Herr lab as UC Berkeley. We are grateful to Lawrence Berkeley National Lab ENG Division and UC Berkeley Biological Imaging Facility.

