## Supplementary Information for "Electromigration of Charged Analytes Through Immiscible Fluids in Multiphasic Electrophoresis"

We introduce $L'$ and $w$ as the axial and radial dispersion characteristic length scales, respectively, with $w$ being the microwell's radius and $L'$ derived from $w$ as follows [1]:

$$L' = 2\left(\frac{\mu_{ii}}{\mu_i}\right) w \text{….(1)}$$

Where $\mu_{ii}$ and $\mu_i$ are the electrophoretic mobilities of BSA in the two mediums present at the interface. For example, in case of negative control medium ii will be gel and medium i will be free solution whereas in case of DropBlot medium i will be oil. We calculate the diffusivity and electrophoretic mobility of BSA in mineral oil using Stokes Einstein equation (mineral oil viscosity: 0.1 Pa.s and BSA hydrodynamic radius: 3.9 nm). For the electrophoretic mobilities in PAGE gel and free solution, we use values that has been reported previously ($\mu_{gel}$: $5.25\times10^{-9}$ $m^2V^{-1}s^{-1}$ and $\mu_{free\ solution}$: $1.58\times10^{-8}$ $m^2V^{-1}s^{-1}$) [2]. Using these characteristic lengths, we can compare the time scale of diffusion ($t_{axial} = L'^2/D_{ii}$ and $t_{radial} = w^2/D_{ii}$), with the timescale for electrophoretic injection ($t_{injection} = L/v$). Where $D_{ii}$ and $v$ are the diffusion coefficient and the average velocity under an applied electric field of BSA in medium ii, respectively. We also define Peclet number (Pe) which compares the characteristic time for diffusion and advection as [1]:

$$Pe = \frac{L'}{L}\frac{wv}{D_{ii}} \text{….(2)}$$

Where $L$ is the separation length. This definition allows us to analyze Pe for different separation lengths. The total dispersion itself is the sum of dispersions arising from injection, radial and axial diffusion:

$$\sigma^2 = \sigma_{inj}^2 + \sigma_{radial}^2 + \sigma_{axial}^2 \text{ ….(3)}$$

Where, $\sigma_{inj}^2$, $\sigma_{radial}^2$ and $\sigma_{axial}^2$ represent the peak variance contributions from injection, radial, and axial diffusion, respectively. The temporal and spatial evolution of a sample peak post-injection can be modeled using Taylor-Aris dispersion. According to the Taylor-Aris dispersion model, the variances for a circular microwell geometry are derived as follows [3–5]:

$$\sigma_{inj}^2 = 0.25\left(\frac{\mu_{ii}}{\mu_i}\right)^2 w^2 \text{….(4)}$$

$$\sigma_{radial}^2 = \frac{1}{X}\left(1 - 2\frac{\mu_{ii}}{\mu_i}\right)\left(w - \sqrt{w^2 - 8D_{ii}t}\right)^2 \text{….(5)}$$

$$\sigma_{axial}^2 = 2D_{ii}t \text{….(6)}$$

Here X is a constant determined by the shape of the input response function (IRF) which contributes to the excess variance. For a Gaussian IRF, X=16 [1].

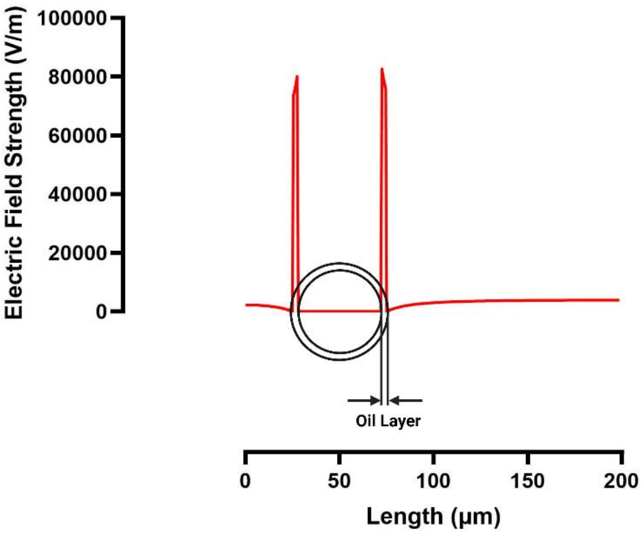


S1: Simulation results of electric field strength across an aqueous droplet, through an oil layer, and into a polyacrylamide gel. The simulation demonstrates a sharp increase in electric field strength upon entering the oil layer, followed by a uniform distribution throughout the gel layer.

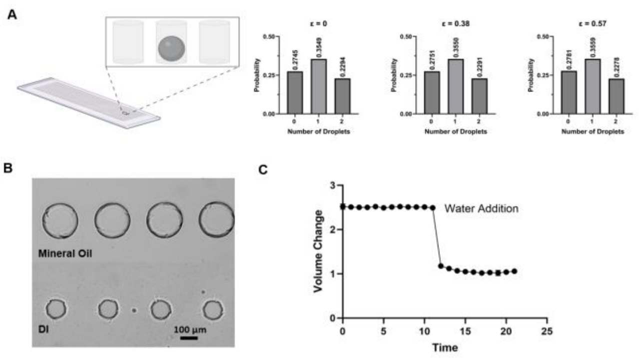


S2: (A) Poisson distribution analysis of 0 1 and 2 droplet settling in microwells with different eccentricities. (B) Microwell expansion and contraction upon oil and water treatment. (C) Microwell hydration dynamics.
